# Bacterial and plant produced lipids can exacerbate the Olive Quick Decline Syndrome caused by *Xylella*

**DOI:** 10.1101/867523

**Authors:** Valeria Scala, Nicoletta Pucci, Manuel Salustri, Vanessa Modesti, Alessia L’Aurora, Marco Scortichini, Marco Zaccaria, Babak Momeni, Massimo Reverberi, Stefania Loreti

## Abstract

*Xylella fastidiosa* is an insect vector-transmitted bacterial plant pathogen associated with severe diseases in a wide range of plants. In last decades, *X. fastidiosa* was detected in several European countries. Among *X. fastidiosa* subspecies, here we study *X. fastidiosa* subsp. *pauca* associated with the Olive Quick Decline Syndrome (OQDS) causing severe losses in Southern Italy. First, we collected *Olea europaea* L. (cv. Ogliarola salentina) samples in groves located in infected zones and uninfected zones. Secondly, the untargeted LC-TOF analysis of the lipid profiles of OQDS positive (+) and negative (-) plants showed a significant clustering of OQDS+ samples apart from OQDS-ones. Thirdly, using HPLC-MS/MS targeted methods and chemometric analysis, we identified a shortlist of 10 lipids significantly different in the infected versus healthy samples. Last, we observed a clear impact on *X. fastidiosa* subsp. *pauca* growth and biofilm formation *in vitro* liquid cultures supplemented with these compounds.

Considering that growth and biofilm formation are primary ways by which *X. fastidiosa* causes disease, our results demonstrate that lipids produced as part of the plant’s immune response can exacerbate the disease. This is reminiscent of an allergic reaction in animal systems, offering the depression of plant immune response as a potential strategy for OQDS treatment.

**Author summary:** Global trade and climate change are re-shaping the distribution map of pandemic pathogens. One major emerging concern is *Xylella fastidiosa*, a tropical bacterium recently introduced into Europe from America. Its impact has been dramatic: in the last 5-years only, Olive Quick Decline Syndrome (OQDS) has caused thousands of 200 years old olive trees to be felled in the southern Italy. *Xylella fastidiosa* through a tight coordination of the adherent biofilm and the planktonic states, invades the host systemically. The planktonic phase is correlated to low cell density and vessel colonization. Increase in cell density triggers a quorum sensing system based on cis 2-enoic fatty acids—diffusible signalling factors (DSF) that promote stickiness and biofilm. Xylem vessels are occluded by the combined effect of bacterial biofilm and plant defences (e.g. tyloses). This study provides novel insight on how *X. fastidiosa* subsp. *pauca* biology relates to the Olive Quick Decline Syndrome. We found that some class of lipids increase their amount in the infected olive tree. These lipid entities, provided to *X. fastidiosa* subsp. *pauca* behave as hormone-like molecules: modulating the dual phase, e.g. planktonic *versus* biofilm. Probably, part of these lipids represents a reaction of the plant to the bacterial contamination.

## Introduction

*Xylella fastidiosa* (Xf) is one of the top 10 plant pathogenic bacteria [1] and is the cause of an environmental emergency within the European Union (EU). Xf has infected a broad host range [2] of plant species; symptoms vary depending on the combination of the host plant and Xf strain [3]. At present *X. fastidiosa* subsp. *pauca*, associated to the Olive Quick Decline Syndrome (OQDS), and three other subspecies, *fastidiosa, multiplex*, and *sandyi* have been identified in Europe [4–7]. Xf subsp. *pauca* was initially introduced from America into Southern Italy and recovered in olive trees [8,9]. This pathogen is associated to the OQDS which has caused losses up to €390 M of the national olive oil production, in the last three years in Italy [10].

Xf is an obligated vector-transmitted pathogen and a xylem-limited bacterium. Inter-host transmission is mediated by xylem sap-feeding insects. The biology of invasion has been well described in the grapevine [11]. From the point of entry in grapevine, Xf moves along the xylem, attaches to its walls and, through a tight coordination of the adherent biofilm and the planktonic states, invades the host systemically [12,13]. The planktonic phase is correlated to low cell density and vessel colonization [14]. Increase in cell density triggers a quorum sensing system based on cis 2-enoic fatty acids—diffusible signalling factors (DSF) that promote stickiness and biofilm [15]. Xylem vessels are occluded by the combined effect of bacterial biofilm and plant tyloses, causing symptoms such as leaf scorch [11,16].

Xf can modulate gene-expression via DSF [17,18]. DSF-mediated quorum sensing determines: a) degradation of the pit membranes to enable cross-vessel diffusion; b) twitching motility of Xf cells; c) adhesion to the xylem surface and biofilm formation. The earlier stage of infection in Pierce disease, consists of evasion by the pathogen of the plant innate immune response and colonization of its vessels, de facto limiting opportunities for reacquisition by the feeding insect vectors [19]. At this stage, Xf is not detected by the plant as a biotic stress, but rather as an abiotic stress (drought and dehydration) [20]. At later stages, the biofilm-based phenotype, consisting of a high density of Xf cells, facilitates reacquisition by the vector and diffusion into other hosts. Only at this point, the plant recognizes the pathogen and mounts an immune response ineffective at preventing Xf colonization and symptoms [21,22]. DSFs act as coordinators of this dual activity of Xf, allowing the switch from the early stage (endophytic lifestyle) into the later stage (insect-acquisition) [11,23]. Lipids appear to be central in the pathogen/host interplay: DSFs trigger the “escape” of the pathogen from the xylem into the vector, ready to change host; contextually, lipidic compounds such as oxylipins, jasmonic acid (JA), free fatty acids, azelaic acid and phosphatidic acid modulate plant defenses acting as signals at the site of infection, as well as in distal parts of the plant [24–26]. In the pathogenic bacteria (e.g. *Pseudomonas aeruginosa*), oxylipins act as hormones for controlling the switch among the different stages of bacterial lifestyle: planktonic, twitching, and biofilm. In *P. aeruginosa*, the oleic acid-derived oxylipins control the virulence in the host and function as autoinducers of a novel quorum sensing system mediating cell-to-cell communication in bacteria [27,28].

In this study, we analyzed the lipidomic profile of a sample of 60 y.o. *Olea europaea* L. cv. Ogliarola salentina symptomatic and symptomless for the OQDS. The profile of 437 lipid compounds was assayed: 186 were found to be differentially accumulated in OQDS positive individuals and 90 were further characterized and quantified by MS/MS spectrometry. Among these, we identified ten compounds that determine the onset of growth and biofilm formation in the pathogen. Importantly, we found that these lipids can be part of plant’s immune response to infection. We thus propose circumscribing plant immunity as a strategy for OQDS treatment.

## RESULTS

### Infected trees exhibit a lipid profile that is distinct from healthy trees

A lipidomic analysis was performed to establish if lipids are differentially accumulated in OQDS symptomatic and symptomless samples. Firstly, the samples were molecularly assayed via real-time PCR to assess the presence or absence of the pathogen and thereon, defined as Xf+ and Xf-. Volcano plot of LC-ToF untargeted analyses on negative-ion scans identified 437 compounds, 186 of which significantly modified (Fold Change >2.0; p <0.05), with 173 upregulated and 13 downregulated (Fig 1). PCA highlighted two primary components (X-axis: 60.48%; Y-axis: 12.05%) that separated the Xf+ and Xf-clusters (Fig 2). Volcano plot analysis on positive-ion scans provided a few significantly modified compounds (S1 Fig).

**Fig 1.**
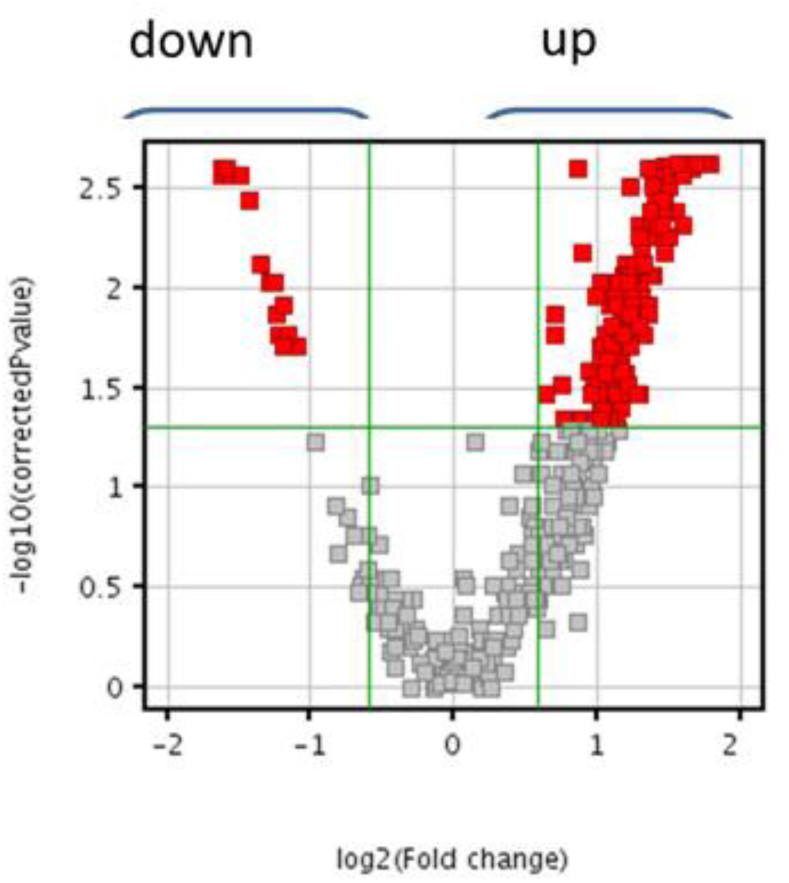
Volcano plot analysis of 437 compounds on negative ion scan, in Xf+ and Xf-. The x-axis shows the fold change to indicate the variation in abundance of compounds present in Xf+ compared to Xf-. The compounds on the right side were more abundant, whereas those on the left side were less abundant in Xf+ conditions. Entities that satisfied the fold change and the *P*-value cut-off of 2.0 and 0.05, respectively, are marked in red.

**Fig 2.**
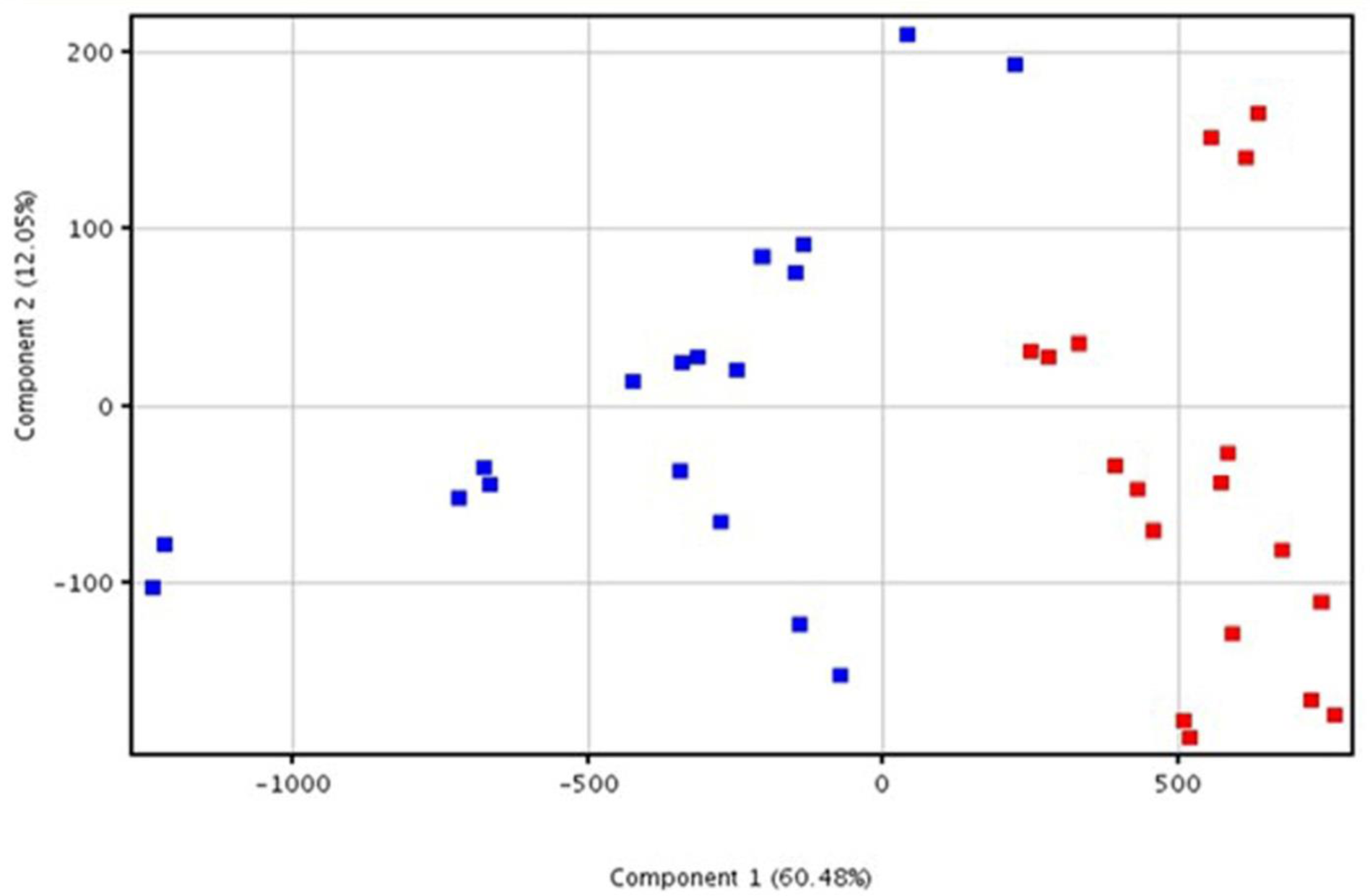
Principal component analysis. Score plot of data generated by HPLC-ECI/TOF-MS analysis of Xf positive (red square) and Xf negative (blue square) samples. The results of the analysis referred to three separate experiments performed in triplicates.

Among the 186 differentially accumulated, 90 compounds were selected and analyzed by a targeted MRM or SIM on the basis of the highest FC and in light of previous results [29] (S1Table). The chemometric analysis highlighted that ten compounds clearly discriminated Xf+ from Xf-samples and were more abundant in Xf+ samples. Namely: seven oleic/linoleic/linolenic acid-deriving oxylipins (9-hydroxyoctadecenoic acid - 9HODE; 9-hydroperoxyoctatrienoic acid - 9HOTrE; 13-hydroxyoctadecenoic acid - 13HODE; 13-hydroperoxyoctatrienoic acid - 13HOTrE; 13-oxo-octadecenoic acid - 13oxoODE; 10-hydroxyoctadecenoic acid −10HODE; 10-hydroperoxyoctamonoenoic acid - 10HpOME); two unsaturated fatty acids (oleic acid - C18:1; linoleic acid - C18:2); and one diacylglycerol [DAG36:4 (18:1/18:3)] (Fig 3 and S1Table).

**Fig 3.**
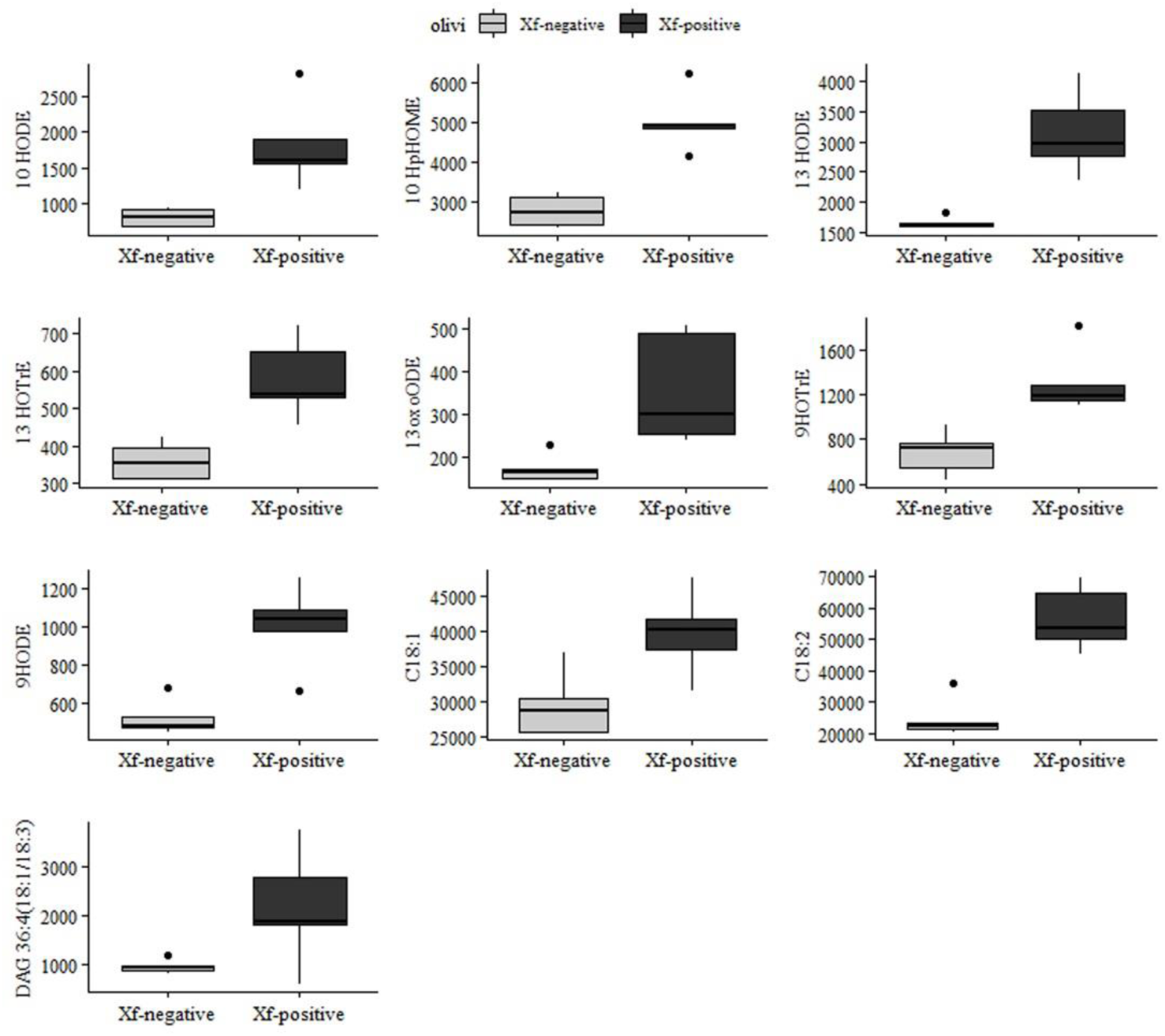
Boxplots of statistically significant compounds analyzed by MRM or SIM. Comparisons between naturally non-infected (Xf negative) and naturally infected (Xf positive) plant samples are displayed on X-axis. The Y-axis shows the normalized relative abundance. Horizontal line in each boxplot indicates the median and black dots represent the outlier samples.

### Bacterial and plant-produced lipids modulate Xf growth and biofilm formation

To elucidate the regulatory role in Xf of the compounds showed in Fig 3, we proceeded to an *in vitro* test of biofilm formation and plancktonic growth. *In vitro* test of biofilm formation indicated that the 7,10-dihydroxyoctamonoenoic acid (7,10-diHOME), the mix 7.10 diHOME and 10-hydroxyoctamonoenoic acid (10-HOME), C18:1 and C18:2 and their diacylglycerols (DAG 36:2 and 36:4), at the concentration of 0.0025 mg ml^−1^, stimulated planktonic growth; at the same concentrations, 9-HODE induced biofilm formation whereas 7,10-diHOME and the mix of 7,10 diHOME and 10-HOME strongly inhibited it (Fig 4). Free fatty acids (linoleic acid, oleic acid) and diacylglicerides (1,3-Dilinoleoyl-rac-glycerol; 1,3-Dioleyl-rac-glycerol), tested at different concentrations, stimulated planktonic growth and did not affect biofilm formation. LOX-derived oxylipins, namely 9HODE, 13HODE, 13OXODE and 9HOTRE promoted biofilm whilst the JA-related 13-HOTRE did not significantly affect it (S2 Fig). The DOX-derived oxylipins, 7,10-DiHOME and 10-HOME, probably produced by the pathogen (as suggested in [13]), strongly inhibited biofilm formation (S3 Fig). Oxylipins did not show an effect on bacterial growth, except for the mixture of 7,10-DiHOME and 10-HOME (S4 Fig).

**Fig 4.**
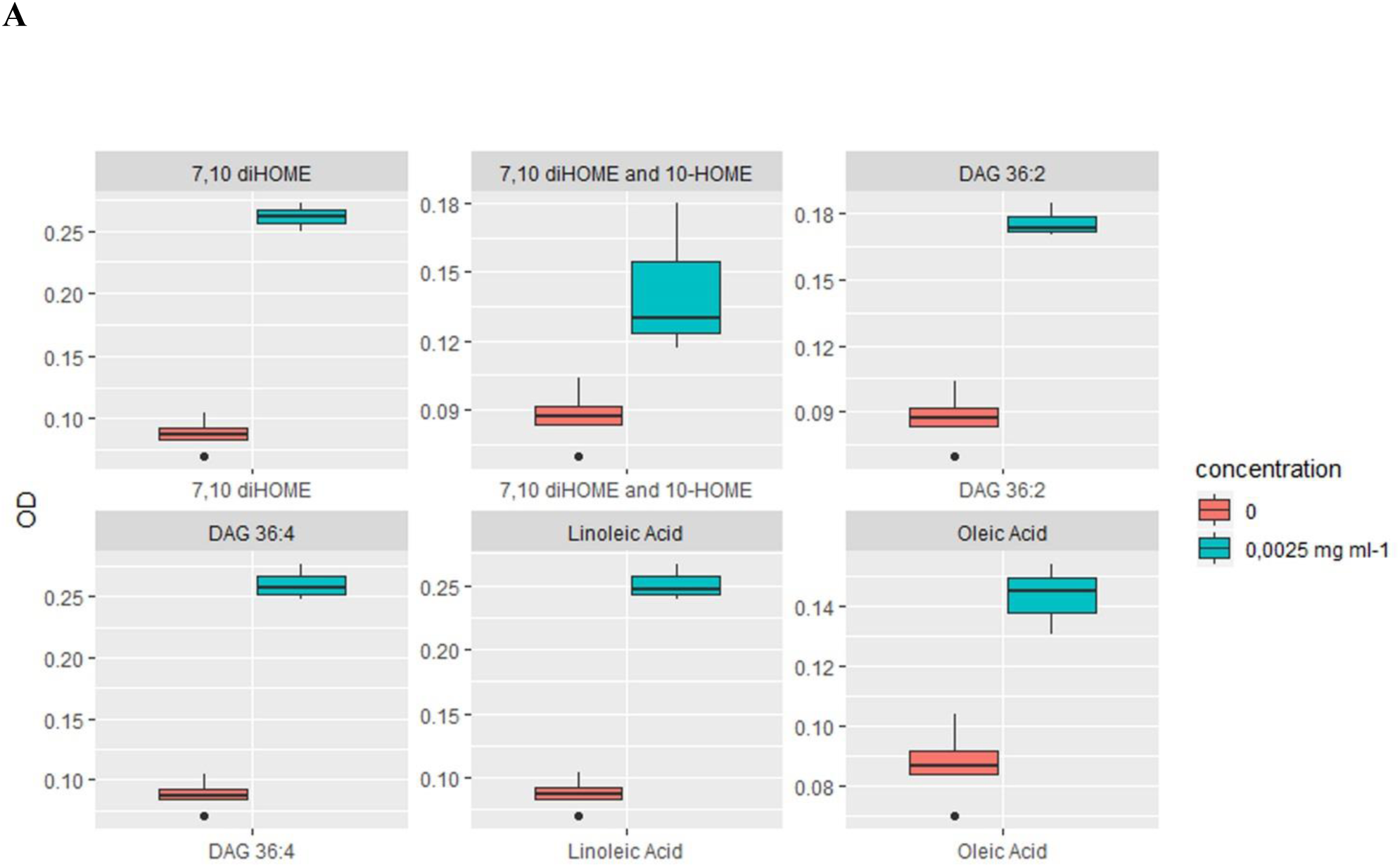
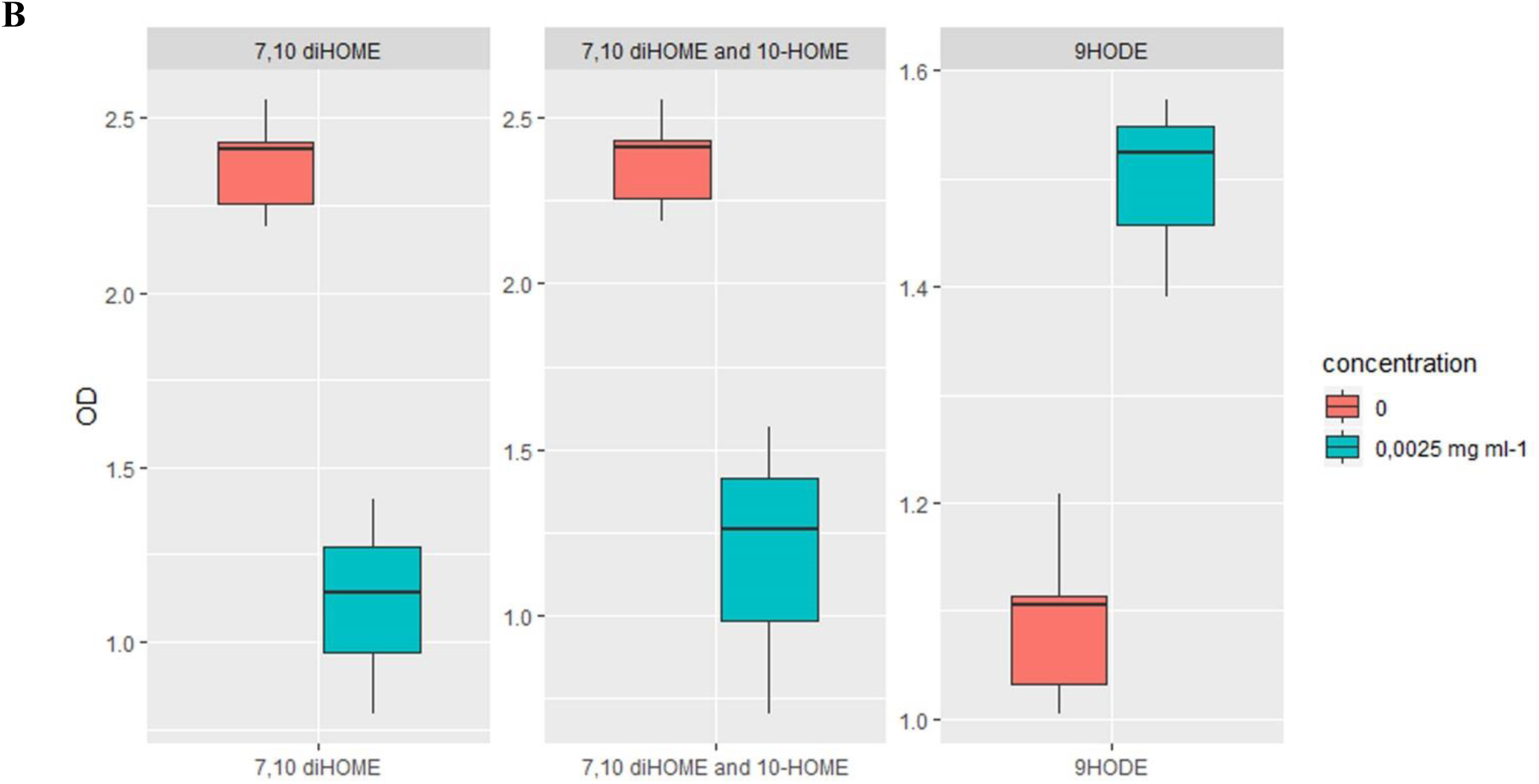
Boxplots of effects caused by lipid compounds on planktonic growth (A) and biofilm formation (B). Selected compounds (FFA, DAG and oxylipins) are the ones with the most relevant effect, measured as OD fold-change, considering 0.0025mg ml^−1^ concentration versus control. The Y-axis displays the optical density (OD). The X-axis displays the compounds (Statistical significance assessed trough Kruskal Wallis test, p-value <0.05). Horizontal line in each boxplot indicates the median and black dots represent the outlier samples.

## DISCUSSION

Lipids are gaining momentum as regulators of host-pathogen interaction; most recently, its role emerged as striking in determining the virulence of an ubiquitary opportunistic bacterial pathogen as *P. aeruginosa* [28]. In other bacterial plant pathogen, such as *Xylella fastidiosa*, other lipids, namely DSF are crucial quorum sensing signals to modulate the pathogen lifestyle [15]. Most information in the biology of *Xylella fastidiosa* referred to one of its subspecies, i.e. *fastidiosa*, and specifically to one of the most important bacterial disease for the grapevine, the Pierce’s disease [21]. In olive trees, there are few reports on the molecular basis of Xf subsp. *pauca* invasion [30,31]. However, in the infection by Xf subsp. *pauca* of *Nicotiana tabacum*, DSFs-like compounds and oxylipins emerged as markers of pathogenic invasion of leaves [29].

Therefore, we focused on the role lipids produced during the host-pathogen interaction might have in the switch between planktonic growth and biofilm formation in Xf subsp. *pauca*: this switch appears to be crucial for xylem occlusion and, therefore, OQDS. We find that the presence of Xf affects the lipidomic profile within the host (Fig 2): specifically, 10 entities clearly discriminate the OQDS+ from OQDS-plants (Fig 3): DAG (36:4), free fatty acids (18:1, 18:2) and LOX- (e.g. 9-HODE) and DOX-derived (e.g. 7,10 diHOME) oxylipins. These lipids may be hallmarks of the plant immune response [32,33] to OQDS, along with bacterial QS-regulation signals [18,28]. We postulate that these entities operate as environment-specific QS signals, or elicitors of QS, since pathogens require exogenous FFA and derived molecules as precursors of the biosynthesis of the auto-inducers to regulate cell-to-cell communication [28]. For instance, DSFs (a miscellaneous of 2-enoic fatty acids) resulting from the coordinated action of the lipase LesA and the crotonase RpfF [18,25], can control the benthonic and planktonic stages of this pathogen.

Our results show that bacterial oxylipins hold back biofilm formation, while LOX-derived oxylipins (e.g. 9-HODE) stimulate it. Since *in silico* analysis (data not shown) did not find any evident LOX-like sequence in the genome of Xf CFBP 8402 strain, it would appear that the messengers responsible for inducing bacterial biofilming are indeed of plant origin. We suggest that the immune response to Xf in olive trees includes LOX-related pathways. This eventually ends up backfiring by causing the pathogen to aggregate and form biofilms that ultimately lead to OQDS—a process resembling an allergic reaction. This is corroborated by evidence of how, in our experiments, the molecular messengers from the pathogen are actually elicitors of the free-moving phenotype, as if Xf would independently not engage in the lifestyle switch to disease-associated biofilm formation. As a side observation, there is evidence of how DAG associated compounds lead to tissue inflammation and damage, in animal systems, through the induction of a disordered immune response [34,35]. The role of plant and bacterial oxylipins (i.e. 13-HODE and 7,10 DiHOME respectively) remains to be clarified; however, we highlight for the first time their pivotal role in the lifestyle of Xf subsp. *pauca* We encourage researchers to investigate oxylipins as targets for the development of much needed new treatments for OQDS environmental emergency.

## MATERIALS AND METHODS

### Study site and sampling procedures

Sampling was carried out in the Apulia region, in October 2017, on 120 individuals of *Olea europaea* L. cv. Ogliarola salentina (60 years old), 60 showing OQDS symptoms (+) and 60 OQDS symptomless (-). The OQDS+ individuals were collected in an olive grove in Copertino in the infected area of the Lecce province (40°16’5.56” N 18°03’15.48” E); the OQDS-individuals were sampled in Grottaglie, Taranto province (40°32’12.98” N 17°26’14.03” E) in an area regarded by the phytosanitary service of the Apulia region as still unaffected by the pathogen. Trees were identified as symptomatic or symptomless following the criteria as previously reported [30]. Pools were generated from equal-weight sub-aliquots of lyophilized samples: six pools from OQDS+ and six from OQDS-. For a comprehensive lipidomic analysis, the pools included the overall analytical complexity of our samples [36,37]. OQDS+ and OQDS-samples were molecularly assayed via real-time PCR [38] and thereon defined as Xf+ and Xf-.

### Lipids analysis

Xylem tissue (0,8 - 1,0 gr) was recovered and lipids extraction and analysis were performed as previously reported [29]. Xf+ and Xf-samples were assayed with the internal reference standards tricosanoic acid, glyceryl tripalmitate d31, and 9-HODEd4. The analysis was carried out at a final concentration of 2µM. The samples were analysed by untargeted lipid analysis conducted with a G6220A TOF-MS, (Agilent Technologies, United States) operating in negative and positive ion scan mode as described in Scala et al. (2018). A sub-group of lipid classes was analysed (fragmentation analysis) by LC-MS/MS (Triple Quadrupole; 6420 Agilent Technologies, United States) as reported [29]; multiple reaction monitoring (MRM) methods were adopted to analyse the most abundant lipid entities (Table S1). MRM data were processed using the Mass Hunter Quantitative software (B.07.00 version). The mass spectrometry analyses were performed three times, each time in technical triplicate (n=9). PCA and significance-fold change analysis (Volcano plot) for untargeted LC-TOF/MS results were performed trough Agilent Mass Analyzer software. Significance tests (T-Student Test, p<0.05) and plots of MRM and SIM results were performed trough R software.

### In vitro assays on planktonic growth and biofilm formation

An *in vitro* test was made to assess the effect of free fatty acids, diacylglycerides, and oxylipins on growth rate and biofilm formation of *Xylella fastidiosa* subsp. *pauca* strain De Donno (CFBP 8402). Xf subsp. *pauca* growth and biofilm formation were evaluated as previously described [39] with some modifications. Briefly, a pure culture of the bacteria was grown for 7 days on PD2 plates, scraped and resuspended in PBS. 10µL of cell suspension (A600 = 0.5 OD) was inoculated in a sterile glass tube containing 1mL of PD2. The free fatty acids (FFA) (Sigma-Aldrich) or diacylglycerides (1,3-Dilinoleoyl-rac-glycerol; 1,3-Diolein) (Sigma-Aldrich) or oxylipins [Cayman chemicals or 7,10 DiHOME and the mix (7,10 DiHOME; 10-HOME) kindly provided by Dr. Eriel Martínez and Javier Campos-Gómez (Southern Research Center, AL, USA] were added to the medium at desired concentrations when required as reported [27]. After 11 days of incubation (28°C; 100 rpm), the total number of cells - planktonic growth (cells in suspension) and biofilm growth (cells adhered to the substrate) was estimated. The lipid compounds were divided into two groups: those dissolved in EtOH (with EtOH) and those dissolved in water (no EtOH). Spectrophotometric absorption of Xf subsp. *pauca* cultures was used to measure growth (600 nm) and biofilm formation (595 nm). For compounds dissolved in EtOH, references with a corresponding concentration of EtOH but without the compound were used as background. Impact on growth / impact on biofilm are defined as the amount of growth/biofilm formation minus the background, normalized to the growth/biofilm formation in a medium in the absence of added EtOH or lipids. A positive impact indicates values more than no-lipid controls (i.e. improved growth or biofilming), whereas a negative impact indicates values less than no-lipid control (i.e. inhibited growth or biofilming). EtOH at similar molarity to lipids was additionally used as a point of reference of how Xf subsp. *pauca* growth and biofilms were affected. The experiments were performed in biological triplicate for each treatment and carried out three times (total repetitions per treatment n=9). Multiple comparison with Kruskal-Wallis test (p value< 0.05) and Fisher’s LSD post-hoc test, with Bonferroni correction, were run on R software to individuate significant (p<0.05) groupings within the different treatments.

## Acknowledgements

The study was funded by MIPAAFT, Project Oli.Di.X.I.It (“OLIvicoltura e Difesa da Xylella fastidiosa e da Insetti vettori in Italia”), D.M. 23773 del 6/09/2017, Project SALVAOLIVI (“Salvaguardia e valorizzazione del patrimonio olivicolo italiano con azioni di ricerca nel settore della difesa fitosanitaria”), D.M. 33437 del 21-12-201 and by Regione Puglia agreement: “Strategie di controllo integrato per il contenimento di Xylella fastidiosa in oliveti pugliesi ed analisi epidemiologiche del complesso del disseccamento rapido dell’olivo (CoDiRO). B.M. and M.Z. were supported by a start-up fund from Boston College and by an Award in Biomedical Excellence from the Smith Family Foundation.

## Supporting information

**S1 Fig. Volcano plot analysis of 524 compounds on positive ion scan, in Xf+ and Xf-.** The x-axis shows the fold change to indicate the variation in abundance of compounds present in Xf+ compared to Xf-. The compounds on the right side were more abundant, whereas those on the left side were less abundant in Xf+ conditions. Entities that satisfied the fold change and the *P*-value cut-off of 1.5 and 0.05, respectively, are marked in red.

**S2 Fig. Impact on biofilm production at different concentrations of individually added oxylipins to *in vitro* cultures of *Xylella fastidiosa* subsp. *pauca* strain De Donno (CFBP 8402)**. Impact is expressed as OD595 fold-change compared to the control. Concentrations are expressed in molarity. The effect of ethanol at equivalent and higher molarities is plotted as a reference.

**S3 Fig. Impact on biofilm production at different concentrations of individually added lipids (DAG 36:2, DAG 36:4, oleic and linoleic acid, 7**,**10-DiHOME and 7**,**10-DiHOME and 10-HOME mix) to *in vitro* cultures of *Xylella fastidiosa* subsp. *pauca* strain De Donno (CFBP 8402).** Impact is expressed as OD595 fold-change compared to the control, the effect of the corresponding amounts of ethanol employed to dissolve the compounds has been subtracted. Concentrations are expressed in molarity. The effect of ethanol at equivalent and higher molarities is plotted as a reference.

**S4 Fig. Impact on growth development at different concentrations of individually added lipids (DAG 36:2, DAG 36:4, oleic and linoleic acids, 7**,**10-DiHOME and 7**,**10-DiHOME and 10-HOME mix) to *in vitro* cultures of *Xylella fastidiosa*.** Impact is expressed as OD600 fold-change compared to the control, the effect of the corresponding amounts of ethanol employed to dissolve the compounds has been subtracted. Concentrations are expressed in molarity. The effect of ethanol at equivalent and higher molarities is plotted as a reference

**S1 Table** A, **B, C. Analyzed lipid entities and relative parameters for Single Ion Monitoring (SIM) and Multiple Reaction Monitoring (MRM) experiments**. a) MRM analysis method for oxylipins; b) MRM method for phospholipids, glycerolipids, ornitholipids, bactophenols; c) SIM method for free fatty acids.

